# AI-Guided Discovery and Optimization of Antimicrobial Peptides Through Species-Aware Language Model

**DOI:** 10.1101/2025.05.20.654992

**Authors:** Daehun Bae, Minsang Kim, Jiwon Seo, Hojung Nam

**Affiliations:** School of Electrical Engineering and Computer Science, Gwangju Institute of Science and Technology (GIST), Buk-gu, Gwangju, 61005, Republic of Korea; Department of Chemistry, Gwangju Institute of Science and Technology (GIST), Buk-gu, Gwangju, 61005, Republic of Korea; AI Graduate School, Gwangju Institute of Science and Technology (GIST), Buk-gu, Gwangju, 61005, Republic of Korea

**Keywords:** Antibiotics, Antimicrobial Peptides, Multi-drug Resistant, AI-based Screening, Protein Language Model

## Abstract

The rise of antibiotic-resistant bacteria drives an urgent need for novel antimicrobial agents. Antimicrobial peptides (AMPs) show promise solutions due to their multiple mechanisms of action and reduced propensity for resistance development. This study introduces LLAMP (Large Language model for AMP activity prediction), a target species-aware AI model that leverages pre-trained language models to predict minimum inhibitory concentration (MIC) values of AMPs. Analysis of attention values allowed us to pinpoint critical amino acid residues (e.g., Trp, Lys, and Phe). Our work demonstrates the potential of AI to expedite the discovery of peptide-based antibiotics to combat antibiotic resistance.

## Introduction

Antibiotic resistance has emerged as one of the most pressing global health challenges, with the World Health Organization (WHO) warning of a potential “post-antibiotic era” where common infections could once again become lethal[1]. The rapid spread of multi-drug resistant (MDR) bacteria has outpaced the development of new antibiotics, emphasizing the urgent need for innovative strategies to discover and develop novel antimicrobial agents[2–4]. In recent years, antimicrobial peptides (AMPs) have attracted considerable attention as potential alternatives to traditional antibiotics[5–7]. Characterized by their multiple mechanisms of action, such as disrupting cell membranes and targeting intracellular nucleic acids and proteins, and their lower likelihood of promoting resistance development, AMPs have garnered significant attention as promising candidates for next-generation antibiotics. However, despite their considerable potential, the discovery of novel AMP lead sequences often requires the high expenses and prolonged timelines inherent in traditional structure-activity relationship (SAR)-based experiments.

Artificial intelligence (AI) has emerged as a transformative force in the face of these challenges, altering the field of antibiotic discovery. The adoption of AI technologies in the drug discovery pipeline has demonstrated the potential to accelerate both the identification and optimization of innovative therapeutic agents[8–10]. In the context of AMP discovery, AI can be employed to predict the antimicrobial activity of peptides, enabling the rapid screening of vast libraries and the identification of promising hit peptides for further experimental validation[11–16]. AI models for AMP discovery are based on the physicochemical properties of peptide molecules as inputs and exploit various architectures, including deep neural networks (DNN), convolutional neural networks (CNN)[17], long short-term memory (LSTM) networks[18], and bidirectional LSTM (Bi-LSTM) networks[19]. Moreover, the recent emergence of language models based on transformer architecture[20] is increasingly being integrated into the AMP field. Protein language models have gained significant traction in AMP research. Models such as ProtTrans, ESM-1b, and ESM-2 have been widely adopted[21–23]. These models excel at processing peptide sequences as input, autonomously learning, and leveraging relevant features. This capability has proven instrumental in facilitating the discovery of novel AMPs, as demonstrated in several recent studies[24–26].

Despite their numerous strengths, previous studies have encountered significant limitations, particularly in their methodology for classifying peptide sequences as either antimicrobial peptides (AMPs) or non-AMPs. Additionally, a notable deficiency in many of these studies is the lack of consideration for the target bacterial species within their predictive models. To address these challenges, we introduce LLAMP (Large Language model for AMP activity prediction), a species-aware activity predictor. LLAMP represents an innovative AI model that employs advanced deep learning techniques, leveraging the power of pre-trained language models. Based on ESM-2, a protein language model utilizing transformer architecture, LLAMP is designed to predict the minimum inhibitory concentration (MIC) values of peptide sequences against different bacterial species. Our approach involved several key steps (**Figure 1a, b**). First, we collected 1.7 million extensive peptide sequence data. Second, we applied ESM-2 for masked language modeling (MLM), resulting in a peptide-tuned version of ESM-2. Next, we encoded genomic information of various bacterial species to represent the target species. Finally, we fine-tuned the peptide-tuned ESM-2 model, integrating the genomic information to create LLAMP. The LLAMP model is capable of predicting MIC values for AMPs based on both the peptide sequence and the target bacterial species. By providing species-aware MIC predictions, LLAMP aims to improve the efficiency of AMP design and screening processes. This is particularly valuable in the fight against newly emerging MDR bacteria, where rapid and targeted antibiotic development is crucial. Moreover, the attention mechanisms within LLAMP provide valuable insights into the significance of specific amino acid residues. Our approach offers a powerful new tool in the ongoing battle against antibiotic resistance.

**Figure 1.**
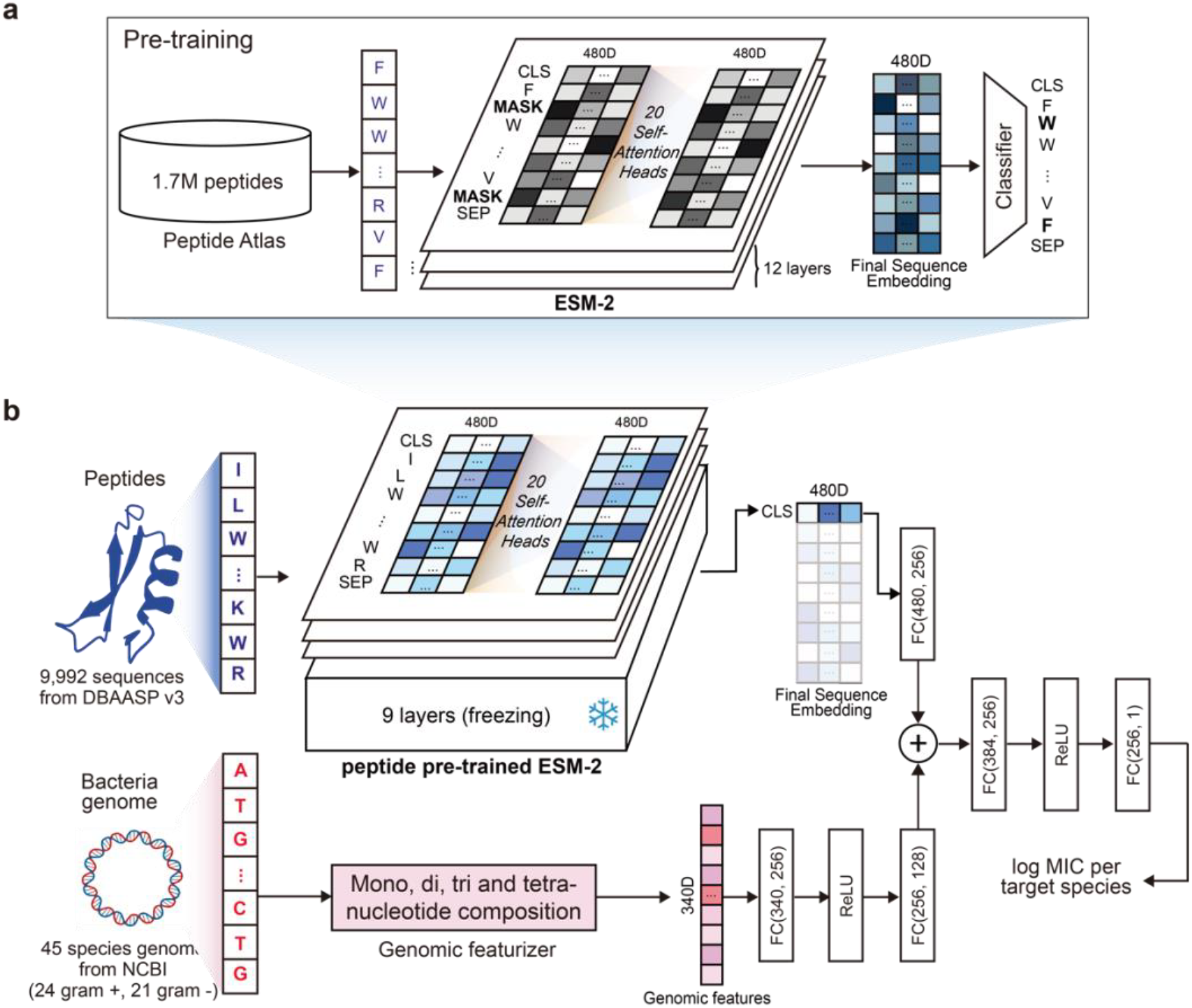
AI-based AMP discovery. (**a**) Pre-training process of the LLAMP model. In the pre-training process, the ESM-2 protein language model is trained with ∼1.7M peptide sequence so that the model can learn sequence patterns of peptides. (**b**) In the fine-tuning process, the LLAMP model learns species-aware activities of peptides and predicts MIC values.

## Results

### LLAMP is a well-trained peptide language model

In this study, we introduce LLAMP, a cutting-edge AI model designed for species-aware antibiotic activity prediction. LLAMP leverages advanced machine learning techniques, including the use of pre-trained language models. At its core is ESM-2[23], a protein language model built on the transformer architecture, which is utilized to predict the minimal inhibitory concentration (MIC) values for peptide sequences targeting specific bacterial species.

The development process began with the collection of peptide sequence data (please see **Datasets** in **Materials and Methods** for more details), on which ESM-2 was applied for masked language modeling (MLM). This process involved masking a portion of the amino acids in each peptide sequence and training the model to predict the masked tokens, leading to the generation of a peptide-optimized version of ESM-2 (peptide-tuned ESM-2).

To verify the proficiency of the peptide-tuned ESM-2 model in interpreting peptide language, we assessed its accuracy in MLM prediction and pseudo-perplexity using a validation set of 69,242 peptide sequences. The accuracy of MLM prediction was evaluated by comparing the model’s top-k predicted amino acids at each masked position with the actual amino acids (**Figure 2a**). The peptide-tuned ESM-2 model demonstrated higher accuracy compared to the original ESM-2 model in predicting the masked amino acids, indicating its ability to capture the underlying patterns and dependencies in peptide sequences.

**Figure 2.**
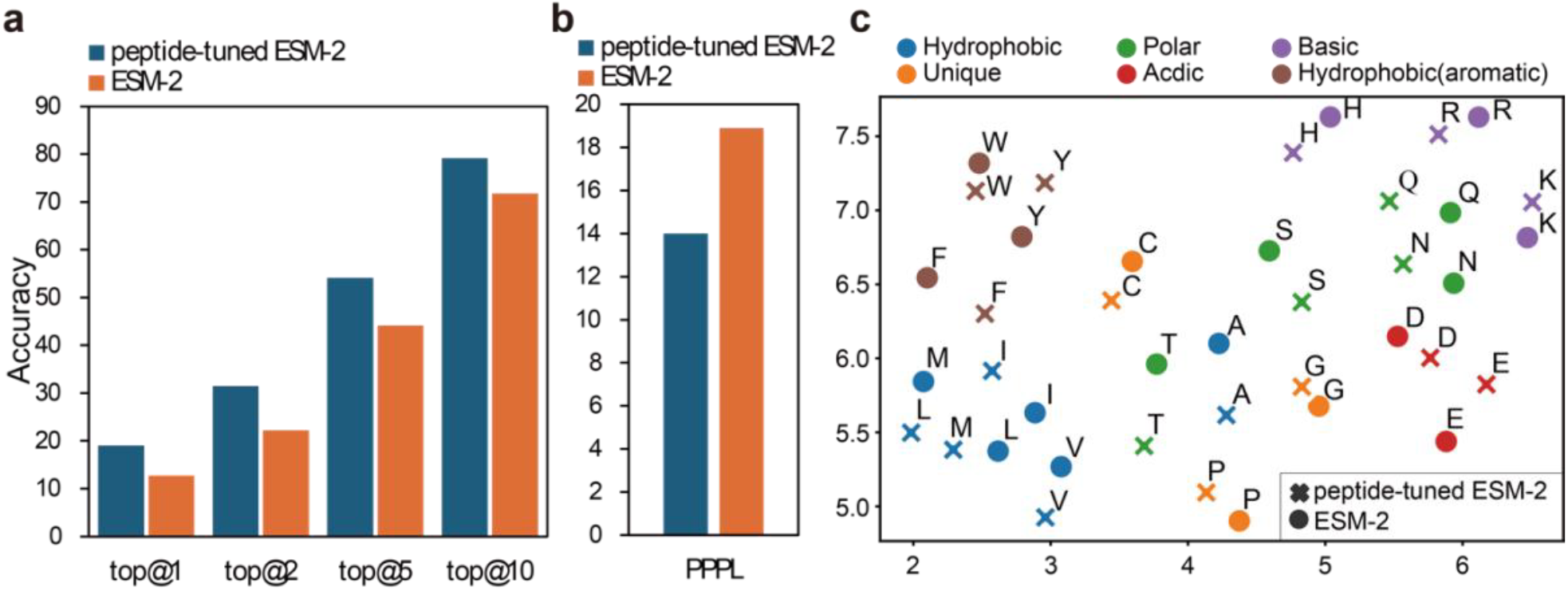
Evaluation of the peptide-tuned ESM-2 model and visualization of amino acid embeddings. (**a**) The accuracy of masked language modeling (MLM) prediction results for the top-*k* most probable amino acids at each masked position. (**b**) Pseudo-perplexity (PPPL) plot comparing the performance of the peptide-tuned ESM-2 model with the original ESM-2 model on a validation set of 69,242 peptide sequences. Lower pseudo-perplexity indicates better performance in capturing the peptide language. (**c**) UMAP visualization of amino acid token embeddings derived from both the original ESM-2 and peptide-tuned ESM-2 models. The peptide-tuned ESM-2 embeddings show distinct clustering based on amino acid properties, demonstrating the model’s improved understanding of peptide language.

**Figure 3.**
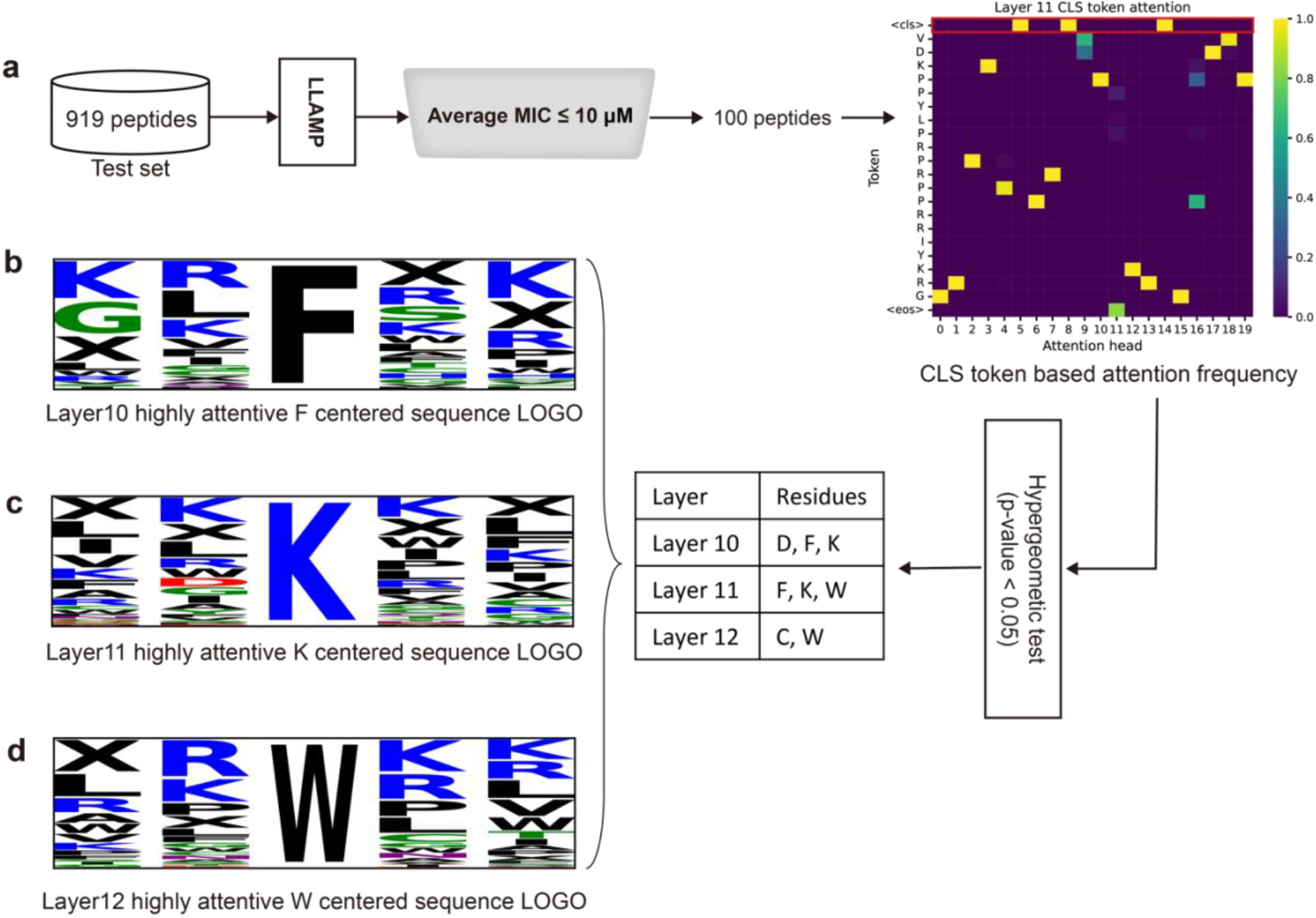
Attention analysis based AMP sequence optimization. (**a**) Whole attention analysis pipeline. In attention analysis, LLAMP CLS token attention is used with 100 active peptides so that the residue-wise frequencies are calculated and important residues are extracted. (**b**), (**c**) and (**d**) are highly attentive F (phenylalanine), K (lysine) and W (tryptophan) centered sequence LOGO for each layer 10, 11 and 12. X residue means terminal padding residue.

Pseudo-perplexity (PPPL), a measure used to evaluate the uncertainty of a language model about a given sequence of text, was also employed to assess the model’s performance[27]. As shown in **Figure 2b**, the pseudo-perplexity of the peptide-tuned ESM-2 model decreased significantly during the training process, reaching its minimum value of 14.0, compared to the initial value of 18.9 for the original ESM-2 model. This reduction in pseudo-perplexity indicates a substantial improvement in the model’s understanding of peptide languages and ability to predict masked residues of peptide sequences after pre-training.

To further investigate the efficacy of pre-training, we analyzed the alterations in amino acid token embeddings, both pre- and post-training, utilizing UMAP visualization[28]. The results, depicted in **Figure 2c**, exhibit a pronounced shift in the embeddings from the baseline ESM-2 model to the peptide-tuned ESM-2. This transition suggests a refined adaptation to peptide-specific languages. Remarkably, the amino acid embeddings demonstrate a tendency to cluster by their respective properties, providing compelling evidence that the peptide pre-training has significantly enhanced the model’s capacity to comprehend and interpret peptide language.

The peptide-tuned ESM-2 model’s enhanced performance in MLM prediction accuracy and reduced pseudo-perplexity demonstrates its effectiveness in capturing the intricate patterns and dependencies within peptide sequences. This optimized model serves as a strong foundation for the subsequent development of the LLAMP framework, enabling more accurate predictions of antimicrobial peptide activity against specific bacterial species.

### LLAMP outperforms in the prediction of species-aware AMP activity

To confirm the predictive performance of the LLAMP model at the species level, we conducted a comprehensive evaluation of its species-aware performance using the fine-tuning data set collected from DBAASP v3 (please see **Datasets** in **Materials and Methods** for more details). The performance of LLAMP was assessed on 35 target species. Notably, 54% of the species exhibited PCC greater than 0.7, indicating a strong predictive performance. However, certain species, such as *Listeria innocua*, showed limited predictive capability, likely due to insufficient training data. This highlights the importance of having a diverse and representative dataset for training the model to ensure robust performance across a wide range of species.

The resulting model, LLAMP, was then compared against five machine learning-based models, two deep learning-based models, and two BERT[29]-based baseline models using the preprocessed test dataset (**Table 1**). The average prediction performance of LLAMP and the baseline models for 35 target species was evaluated using a test dataset. As presented in **Table 2**, LLAMP demonstrated superior performance across all metrics, including Pearson correlation coefficient (PCC) of 0.735, coefficient of determination (R^2^) of 0.536, mean absolute error (MAE) of 0.379, mean squared error (MSE) of 0.272, and root mean squared error (RMSE) of 0.521. These results highlight the overall effectiveness of LLAMP in accurately predicting MIC values for AMPs based on their peptide sequences, outperforming the baseline models.

**Table 1.** Dataset Statistics for training, validation, and testing in peptide MLM tuning and regression fine-tuning.

| Dataset | Origin | Split type | # Peptides | # Species | # Entities |
| --- | --- | --- | --- | --- | --- |
| Pre-training | PeptideAtlas[30] | Train | 1,661,800 | - | - |
|  |  | Validation | 69,242 | - | - |
|  |  | Total | 1,731,042 | - | - |
| Fine-tuning | DBAASP v3[32] | Train | 7,344 | 35 | 29,394 |
|  |  | Validation | 918 | 35 | 3,683 |
|  |  | Test | 919 | 35 | 3,741 |
|  |  | Both-unseen | 811 | 10 | 982 |
|  |  | Total | 9,992 | 45 | 37,800 |

**Table 2.** Prediction performances on the test and both both-unseen sets.

| Set | Type | Method | PCC | $R^2$ | MAE | MSE | RMSE |
| --- | --- | --- | --- | --- | --- | --- | --- |
| Test set | Machine learning-based architectures | RF[38] + CTD[39] | 0.709 | 0.491 | 0.423 | 0.298 | 0.546 |
|  |  | XGB[40] + CTD | 0.651 | 0.407 | 0.441 | 0.347 | 0.589 |
|  |  | SVM[41] + CTD | 0.627 | 0.390 | 0.454 | 0.357 | 0.600 |
|  |  | RF + ESM-2 | 0.680 | 0.443 | 0.448 | 0.326 | 0.571 |
|  |  | XGB + ESM-2 | 0.644 | 0.407 | 0.452 | 0.347 | 0.589 |
|  | Deep learning-based architectures | CTD + FC | 0.581 | 0.317 | 0.495 | 0.400 | 0.633 |
|  |  | Embedding + CNN | 0.680 | 0.451 | 0.420 | 0.322 | 0.567 |
|  | BERT-based architecture | protBERT-BFD[21] | 0.644 | 0.390 | 0.443 | 0.358 | 0.598 |
|  |  | ESM-2 | 0.723 | 0.523 | 0.388 | 0.280 | 0.529 |
|  |  | LLAMP (proposed) | **0.735** | **0.536** | **0.379** | **0.272** | **0.521** |
| Both-unseen set | Machine learning-based architectures | RF + CTD | 0.640 | 0.402 | 0.458 | 0.352 | 0.593 |
|  |  | XGB + CTD | 0.608 | 0.335 | 0.475 | 0.391 | 0.626 |
|  |  | SVM + CTD | 0.480 | 0.180 | 0.539 | 0.482 | 0.695 |
|  |  | RF + ESM-2 | 0.650 | 0.415 | 0.443 | 0.344 | 0.587 |
|  |  | XGB + ESM-2 | 0.577 | 0.273 | 0.487 | 0.428 | 0.654 |
|  | Deep learning-based architectures | CTD + FC | 0.541 | 0.243 | 0.511 | 0.445 | 0.667 |
|  |  | Embedding + CNN | 0.604 | 0.300 | 0.487 | 0.412 | 0.642 |
|  | BERT-based architecture | protBERT-BFD | 0.646 | 0.363 | 0.457 | 0.375 | 0.612 |
|  |  | ESM-2 | 0.618 | 0.337 | 0.469 | 0.390 | 0.625 |
|  |  | LLAMP (proposed) | **0.668** | **0.418** | **0.428** | **0.342** | **0.585** |

To further investigate the model’s performance under more challenging conditions, we employed a both-unseen dataset, which consists of peptide sequences and target species that were not encountered during the training process. This dataset allows for a rigorous assessment of the model’s generalizability and ability to handle novel data. Remarkably, 50% of these species exhibited a PCC score exceeding 0.7, reinforcing our confidence in the model’s capability to accurately predict AMP activity against novel species that were not part of the training data. These outcomes demonstrate the robustness of LLAMP across a variety of species and highlight its potential utility in identifying active AMPs effective against newly emerged or antibiotic-resistant species. Also, as shown in **Table 2**, LLAMP exhibited superior performance compared to the baseline models on this both-unseen dataset, with a PCC of 0.668, R^2^ of 0.418, MAE of 0.428, MSE of 0.342, and RMSE of 0.585.

Lastly, to further validate LLAMP’s performance, we compared it to previous models that predict MICs for specific target species^13-14^, using datasets sourced from their respective works. As shown in **Table 3**, when training, validating, and testing using the ESKAPEE dataset, LLAMP demonstrated similar or better performance on the test set compared to the other models. It is important to note that while both LLAMP and ESKAPEE-MICpred can predict species-aware AMP activity, ESKAPEE-MICpred is only able to support seven species and lacks the generalization ability to predict activity for new species. In contrast, LLAMP, trained on a broader range of species, exhibited a stronger predictive ability for new species, highlighting its greater versatility and generalizability. Furthermore, LLAMP showed a notable improvement in performance on the GRAMPA-*E*.*coli* set, with a PCC increase from 0.770 to 0.811 and a decrease in RMSE from 0.501 to 0.454. This improvement highlights the effectiveness of LLAMP in accurately predicting AMP activity against specific bacterial species, even when compared to models designed explicitly for those species.

**Table 3.**
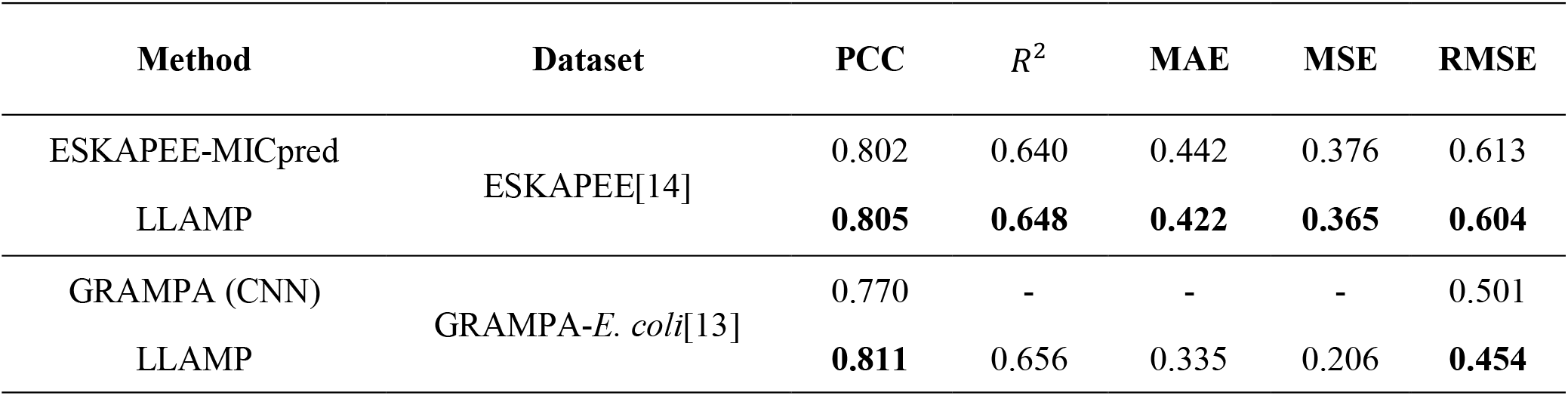
Performance of our model and previous studies.

### The ablation study unveils the critical roles of pre-training and fine-tuning

To identify the key factors contributing to the model’s enhanced performance, we conducted an ablation study by testing the model under various configurations. These cases included (i) without peptide pre-training, (ii) with all transformer encoder layers frozen (all freeze), and (iii) with a reduced number of species in training.

In the “without peptide pre-training” scenario, we used the original ESM-2 model directly for fine-tuning, without any peptide-specific pre-training. The “all freeze” scenario involved freezing the transformer encoder layers during fine-tuning, while the “reduced species” scenario entailed training with only eight species. We evaluated the results of these ablation experiments on the test set (cases 1 and 2) and the both-unseen set (cases 1, 2, and 3).

The “all freeze” scenario led to a substantial decrease in performance on both sets, highlighting the crucial role of fine-tuning the transformer encoder layers. Fine-tuning these layers appears to stabilize the model and facilitate its specialization in regression tasks, enabling it to capture the nuances and complexities of the antimicrobial activity prediction problem. Conversely, the absence of peptide pre-training resulted in a slight performance drop, suggesting that peptide pre-training plays a valuable role in enhancing the model’s understanding of peptide languages and their inherent properties. Lastly, our analysis revealed that decreasing the number of species used in training from 35 to 8 led to worse performance on the bot-unseen set. This finding highlights the importance of a diverse training set in improving the model’s generalization ability. By exposing the model to a wider range of species during training, it can learn to capture the variability and complexity of antimicrobial activity across different bacterial species, resulting in more robust and accurate predictions.

## Discussion

Our study demonstrates the successful development and application of LLAMP, a species-aware AI model that advances the field of antimicrobial peptide discovery in several key aspects. First, LLAMP’s superior performance compared to existing prediction models (PCC of 0.735 and R^2^ of 0.536 on the test set) validates our approach of combining peptide pre-training with species-specific genomic features. Particularly noteworthy is LLAMP’s ability to generalize to previously unseen bacterial species, achieving a PCC of 0.668 on the both-unseen dataset. This capability is crucial for addressing emerging pathogens and MDR bacteria.

While our results are promising, several areas warrant further investigation. The variation in prediction accuracy across different bacterial species suggests room for improvement in species representation during training. Additionally, the development of resistance and *in vivo* efficacy of the identified peptides needs to be evaluated in future studies. Looking ahead, LLAMP’s framework could be extended to predict activity against other pathogens, including fungi and viruses, and to optimize other peptide properties such as stability and cell penetration. The integration of additional structural and physicochemical features could further enhance prediction accuracy and guide peptide optimization.

This work establishes a powerful platform for accelerating the discovery and optimization of antimicrobial peptides, offering a promising approach to address the growing challenge of antibiotic resistance. The combination of AI-driven prediction, rational design, and experimental validation provides a robust framework for developing next-generation antimicrobial therapeutics.

## Materials and Methods

### Datasets

The datasets utilized in this study are detailed in **Table 1**. Our model consists of two parts: peptide MLM tuning and regression fine-tuning. First, in the peptide MLM tuning part, the context of the peptide is learned by inputting the sequence information of the peptide. The dataset for this phase was sourced from the PeptideAtlas database (12/2022), comprising 5.5 million peptide sequences from various organisms[30]. To eliminate redundant sequences, we employed the CD-HIT program[31] with a 90% threshold, yielding 1,731,042 unique peptide sequences. Of these, 96% were allocated for training and the remaining 4% for validation.

Next, in the regression fine-tuning for the species-aware activity prediction model, we used DBAASP v3[32] and reference genomes of bacteria obtained from NCBI[33]. The DBAASP v3 database, a manually curated resource, offers activity data for various bacterial species, including minimum inhibitory concentration (MIC) values for 16,968 monomer peptide sequences targeting specific bacteria. We curated the database based on multiple criteria.

Consequently, the refined dataset for fine-tuning comprises 9,992 peptide sequences across 45 species. We allocated 35 of these species to the training, validation, and test sets, distributing them in an 8:1:1 ratio based on the number of peptides. Moreover, 10 species were reserved for an external set.

Furthermore, we incorporated and utilized the ESKAPEE-MICpred dataset[14] and the GRAMPA (*E. coli*) dataset[13] to ensure a fair comparison of performance.

### Genomic features

To train the species-specific activity prediction model, we extracted features of bacterial species from the whole genome sequence downloaded from NCBI Genome (downloaded December 2022)[33]. For feature extraction, the mono(4), di-(16), tri-(64), and tetra-(256) nucleotide compositions of the genome were obtained and normalized, respectively using Biopython[34,35].

### Protein language model and peptide MLM tuning

ESM-2, inspired by BERT[29], is the state-of-the art (SOTA) model among protein language models, which showed the best performance in contact prediction, and structure prediction of proteins[23]. It offers a range of model sizes, varying from 8 million to 15 billion parameters, and is readily accessible on HuggingFace[36], facilitating easy fine-tuning. For our study, we utilized the ‘esm2_t12_35M-UR50D’ as a baseline model, which features 12 transformer encoder layers and 35 million parameters.

To enhance the protein language model’s proficiency in interpreting peptide language, we further pre-trained ESM-2 using shorter peptide sequences compared to full-length proteins (**Figure 1a)**. Our training process involved randomly masking 15% of the sequences. Within these masked sequences, 80% of the tokens were masked, 10% were replaced with random tokens, and the remaining 10% were left unmasked. This training technique is in line with the pre-training approaches typically used for protein language models.

### Regression fine-tuning

The structure of LLAMP is shown in **Figure 1b**. The LLAMP model takes a pair consisting of a peptide sequence and a bacterial genome as input. The peptide sequence passes through peptide-tuned ESM-2, generating an embedding that represents the sequence. Concurrently, the bacterial genome is transformed into a 340-dimensional vector using a genomic featurizer. This genomic representation is then processed through a series of linear layers before being merged with the peptide embedding. Finally, after the final layer, the MIC value is predicted. For this, the Adam optimizer[37] and MSE loss were used.

### Evaluation metrics

To evaluate the pre-training performance, we utilized pseudo-perplexity, a metric that gauges the model’s understanding of peptide sequences. Pseudo-perplexity, as proposed by Salazar et al.[27], is the exponential of the negative pseudo-log-likelihood of a sequence. Pseudo-perplexity is defined by the following formulas:

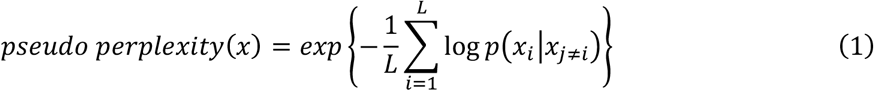

*L* is the length of the input sequence.

Next, to evaluate the performance of the fine-tuned model, we utilized the Pearson correlation coefficient (PCC), R squared (R^2^), mean absolute error (MAE), mean squared error (MSE), and root mean squared error (RMSE) which are standard metrics in regression tasks. The five metrics are defined by the following formulas:

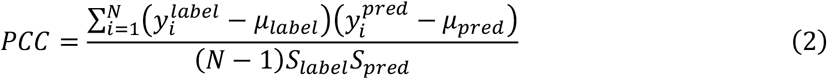

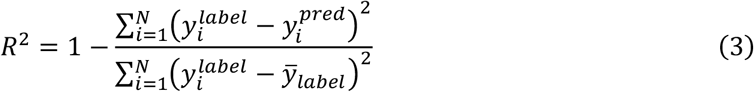

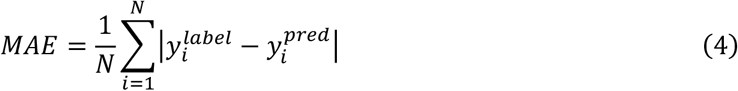

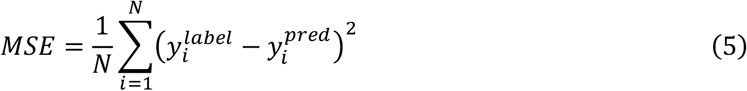

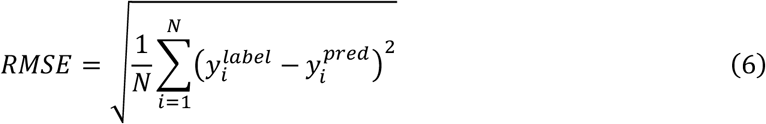

### Attention analysis and AMP sequence optimization

To gain insight into the importance of specific amino acid residues, we analyzed sequences from the fine-tuning test set. We performed attention analysis on the top 100 sequences with the highest average predictive values out of 919 test sequences. The average predicted log10 MIC values for these top 100 sequences ranged from -1.14 to 0.49.

We then input these 100 sequences into the LLAMP model to obtain the attention weights of the CLS token across all heads for each of the three fine-tuned layers. For each layer, we identified residues with attention values exceeding 0.5 and applied a hypergeometric test to each amino acid residue. This analysis allowed us to determine which amino acids were statistically overrepresented among the high-attention residues, providing insights into the model’s focus during prediction.

## Data Availability

The raw datasets are all publicly available.

DBAASP version 3: https://dbaasp.org/PeptideAtlas: https://peptideatlas.org/

GRAMPA: https://github.com/zswitten/Antimicrobial-Peptides/blob/master/data/grampa.csv

ESKAPEE-MICpred: https://eskapee-micpred.anvil.app/

## Author Contributions

D.B., M.K., J.S., H.N. conceptualized the study. D.B. implemented the AI model pipeline. M.K. performed *in vitro* validations. D.B., M.K., J.S., H.N. analyzed and interpreted the results. D.B., M.K. wrote the initial manuscript. J.S., H.N. reviewed and revised the manuscript.

## Acknowledgements

We appreciate the high-performance GPU computing support of HPC-AI Open Infrastructure via GIST SCENT.

